# Dynamic network features of functional and structural brain networks support visual working memory in aging adults

**DOI:** 10.1101/2024.07.30.605891

**Authors:** Josh Neudorf, Kelly Shen, Anthony R. McIntosh

**Affiliations:** Institute for Neuroscience and Neurotechnology, Simon Fraser University, Burnaby, Canada; Department of Biomedical Physiology and Kinesiology, Faculty of Science, Simon Fraser University, Burnaby, Canada

**Author notes:** Corresponding author (ARM).

**Keywords:** resting-state functional magnetic resonance imaging, dynamic functional connectivity, diffusion-weighted magnetic resonance imaging, network control theory, healthy aging, visual working memory

## Abstract

In this work, we investigated the relationship between structural connectivity and the dynamics of functional connectivity and how this relationship changes with age to benefit cognitive functions. Visual working memory (VWM) is an important brain function that allows us to maintain a mental representation of the world around us, but its capacity and precision peaks by around 20 years old and decreases steadily throughout the rest of our lives. This research examined the functional brain network dynamics associated with VWM throughout the lifespan and found that Default Mode Network and Fronto-Parietal Network states were more well represented in individuals with better VWM. Furthermore, transitions between the Visual/Somatomotor Network state and the Attention Network state were more well-represented in older adults, and a network control theory simulation demonstrated that structural connectivity differences supporting this transition were associated with better VWM, especially in middle-aged individuals. The structural connectivity of regions from all states was important for supporting this transition in younger adults, while regions within the Visual/Somatomotor and Attention Network states were more important in older adults. These findings demonstrate that structural connectivity supports flexible, functional dynamics that allow for better VWM with age and may lead to important interventions to uphold healthy VWM throughout the lifespan.

---

The healthy brain constantly adapts to changing contexts and task demands, and even during rest. This adaptation can be described as a fluid motion through a high dimensional space of possible brain states. Of particular interest is the question of how the transitions between one brain state and another may be unique across individuals and tell us something about that person. How is a person’s unique pattern of transitions between brain states related to the brain’s structure, and other features that set individuals apart (e.g., age and cognitive ability)?

These brain states, separable across time and relying on different combinations of brain regions, can be extracted from brain activation signals using modeling methods including Hidden Markov Modelling (HMM; Haussler et al., 1992) and Leading Eigenvector Dynamics Analysis (LEiDA; Cabral et al., 2017). HMM was initially developed for applications to protein and DNA sequencing, and has been used widely for modelling data that follows a sequence, while LEiDA, developed for neuroimaging data, accomplishes similar goals while integrating a step that extracts the leading eigenvector from time × time phase coherence matrices, capturing dominant connectivity patterns while diminishing the effects of noise. Both of these approaches have been applied to neuroimaging data revealing distinct brain states at different points in time relying on separable combinations of regions (e.g., Cabral et al., 2017; S. E. Faber et al., 2024; S. E. M. Faber et al., 2023; Vidaurre et al., 2017).

Important questions arise from these findings that the brain coordinates the function of different combinations of regions depending on when you observe its activity. Namely, how can the brain produce multiple different repertoires of functional synchronization when the structural connections (white matter tracts of axon bundles) between regions are static at short timescales? This question has been investigated previously (e.g., Deco et al., 2011; Honey et al., 2009), and recent research using network control theory (NCT) has demonstrated that the structural connectivity network of the brain is organized to afford transitions between states via efficient stimulation to certain regions of the brain that produce cascading signals and push brain activity into new states (Gu et al., 2015, 2017; Kim & Bassett, 2020; Lynn & Bassett, 2019; Parkes et al., 2023). This work has demonstrated that although brain function can vary widely over time, the structural connectivity does constrain to what extent certain functional states are possible.

On the one hand, NCT has allowed for interesting simulation-based predictions about how structural connectivity constrains network dynamics, and on the other hand, a number of modelling approaches including HMM and LEiDA allow for the data-driven extraction of network states from real-world time series data such as resting-state fMRI. However, the comparison of these structural and functional perspectives of brain dynamics remains a needed research direction. Furthermore, investigating how the dynamics of the structural and functional brain networks differ across stages of adulthood, and how these differences either support or hinder the brain’s ability to perform important cognitive tasks, will give us a better sense of which features of structural and functional brain dynamics are beneficial as opposed to detrimental.

Visual working memory (VWM) represents one of the earliest aspects of cognitive ability to decline over a lifespan, with decreasing capacity and precision starting at around 20 years of age that continues through middle and old age. In fact, by middle age VWM ability is indistinguishable from that of 8 to 9 year olds (Brockmole & Logie, 2013). For this reason, understanding how VWM is supported by the brain could benefit all adults regardless of their current stage in their lifespan. Understanding how brain adaptations stave off this decline could lead to interventions that support these adaptations as early as the beginning of adulthood and have long-term benefits to cognitive health later in life.

VWM allows individuals to remember and mentally manipulate visual information over short time scales, and as such represents a foundational building block for more complex cognitive tasks relying on a mental representation of the world around us, including navigation and spatial problem solving. VWM utilizes multiple systems in the brain important for visual processing of color and shape information, attentional orienting, and working memory encoding, maintenance, and retrieval. Working memory (WM) in general engages a frontoparietal network of brain regions including dorsolateral prefrontal cortex, posterior parietal cortex, and presupplementary motor areas (D’Esposito, 2007; Smith & Jonides, 1999; Wager & Smith, 2003). VWM specifically relies on visual processing regions of the occipital cortex (Harrison & Tong, 2009), and the posterior parietal cortex has been associated with spatial reasoning and attentional processing, becoming more active as the number of items maintained increases to the individual’s WM limit (Todd & Marois, 2004; see Schurgin, 2018 for a review). Dynamic network analyses that consider how different brain networks are utilized over time have demonstrated that WM tasks reduce the overall modularity in the brain compared to rest, resulting in brain networks that communicate with one another more, leading to a whole-brain effort to perform these tasks, and WM training produces more segregated default mode and task positive (i.e., dorsal attention) networks (Bassett et al., 2015; Finc et al., 2020). Network control theory analyses of brain function during a WM task have identified that signalling between the salience network (i.e., ventral attention network), frontoparietal network, and default mode networks predicts task performance, with the anterior insula and dorsolateral prefrontal cortex acting as important regions facilitating this signalling (Cai et al., 2021). Further understanding of the brain network functional dynamics and structural connections supporting these dynamics is needed, especially in the context of how specific subnetworks change the way they support VWM across the lifespan.

Although age is commonly associated with declining cognitive ability, and negative brain changes, we have estimated complexity with multiscale entropy to demonstrate that some changes in the complexity of function activity with age are associated with spared cognitive ability (Heisz et al., 2015), and that different aspects of structural brain network reorganizations were associated with both declining as well as spared cognitive ability in older adults (Neudorf et al., 2024). In particular, increased local interhemispheric connections and specific regional differences in the organization of hub regions were associated with spared cognitive ability (Neudorf et al., 2024).

For the current work, we investigated the dynamic connectivity patterns (brain states) from both functional and structural brain network perspectives, how the time spent in these states and the pattern of transitions between states differ across the lifespan from younger to older adulthood, and whether some of these differences contribute to better VWM in older adulthood. Recent research has demonstrated strong coupling between structural and functional networks in the brain (Benkarim et al., 2022; Neudorf et al., 2022; Sarwar et al., 2021; Schirner et al., 2018), that this coupling is altered across the lifespan (Zimmermann et al., 2016), and that the structural network constrains the range of functional dynamics possible with an individual’s brain network (Gu et al., 2015; Lynn & Bassett, 2019). For this reason, we will examine how differences in structural connectivity with age may support specific functional dynamics to support better VWM.

## Methods

Data came from the Cambridge Centre for Ageing and Neuroscience (Cam-CAN; Shafto et al., 2014) dataset. Data collection followed the Helsinki Declaration, and was approved by the local ethics committee, Cambridgeshire 2 Research Ethics Committee (reference: 10/H0308/50). The full sample of subjects with neuroimaging data included 653 subjects. Participant ages ranged from 18.5 to 88.92 (mean = 54.825, *SD* = 18.593). Younger adult ages ranged from 18.5 to 49.92 (mean = 36.420, *SD* = 8.495, *N* = 279, 146 female, 133 male). Older adult ages ranged from 50.17 to 88.92 (mean = 68.593, *SD* = 10.338, *N* = 373, 184 female, 189 male). A single participant was missing age information. The resting-state functional MRI (rs-fMRI) subsample included 197 subjects that passed our quality control criteria (see below). Participant ages ranged from 18.50 to 88.92 (mean = 48.310, *SD* = 17.388). Younger adult ages ranged from 18.5 to 49.83 (mean = 35.471, *SD* = 8.176, *N* = 114, 67 female, 47 male). Older adult ages ranged from 51.92 to 86.08 (mean = 65.945, *SD* = 9.308, *N* = 83, 43 female, 40 male). The diffusion-weighted MRI (dMRI) subsample included 594 subjects that passed our quality control criteria (see below). Participant ages ranged from 18.50 to 88.92 (mean = 55.414, *SD* = 18.090). Younger adults (YA; age < 50) ranged from 18.50 to 49.92 years (mean = 36.966, *SD* = 8.385, *N* = 244, 131 female, 113 male) and older adults (OA; age > 50) ranged from 50.17 to 88.92 years (mean = 68.275, *SD* = 10.163, *N* = 350, 170 female, 180 male).

### Structural MRI

The T1-weighted Magnetization Prepared RApid Gradient Echo (MPRAGE) sequence was performed using a repetition time (TR) of 2250 ms and echo time (TE) of 2.99 ms, with a flip angle of 9°, field of view (FOV) of 256×240×192mm, and 1×1×1 mm voxel size. The T2-weighted sampling perfection with application-optimized contrasts using different flip angle evolution (SPACE) sequence was performed using a TR of 2800 ms, a TE of 408 ms, a FOV of 256×256×192mm, and 1×1×1 mm voxel size.

### Functional MRI

The rs-fMRI sequence was performed using a Gradient-Echo Echo-Planar Imaging (EPI) sequence with a TR of 1970 ms, TE of 30 ms, flip angle of 78°, FOV of 192×192mm, and 3×3×4.44mm voxel size. This sequence acquired a total of 261 volumes over 8 minutes and 40 seconds, with each volume containing 32 axial slices. The preprocessing of these data relied on the TheVirtualBrain-UK Biobank pipeline (Frazier-Logue et al., 2022), which has updated the FMRIB Software Library (*FSL*; Jenkinson et al., 2012) based UK Biobank pipeline (Littlejohns et al., 2020) to account for issues that can occur due to atrophy in aging brains using quality control methods to minimize artifacts (Lutkenhoff et al., 2014). This pipeline also outputs parcellation-based blood oxygen level dependent (BOLD) time-series data which were used for the dynamic functional connectivity analyses.

Using 41 imaging-derived phenotypes (IDPs) related to the T1w and T2w structural image quality, rs-fMRI imaging quality, and structural-functional registration from the TVB UKBB pipeline (Frazier-Logue et al., 2022) as predictor variables and human rated scores of rs-fMRI quality (based on visual assessment of the fMRI fieldmaps, motion, registration, *FSL MELODIC* independent component labelling accuracy, functional connectivity matrix, and timeseries carpet plot) on a scale of 1-5 (1 is excellent, 2 is good, 4 is poor, and 5 is very poor) as the criterion variable, a random forest regression machine learning approach was trained to predict rs-fMRI quality on a subset of the Cam-CAN data (140 participants). The *auto-sklearn* (Feurer et al., 2015) Python library was used to aid selection the best performing machine learning algorithm and parameters, and Random Forest Regression was selected (*scikit-learn*; Pedregosa et al., 2011). Human rated quality was rescaled to the range of 0 to 1, with an original score of 1 corresponding to 0.1 and with 0.2 increments between scores. When applying this model to the unrated subjects (N=499), a score less than 0.4 corresponded to a passing value, a score greater than 0.6 corresponded to a failing value, and scores between 0.4 and 0.6 were selected for manual human rating. Using this same procedure with the manually human rated subjects (N=140), in a K-fold validation scheme (K=5) repeated over 100 iterations, setting aside the subjects in the medium range of 0.4 to 0.6 and looking only at those subjects identified confidently as good or bad we observed a false negative detection of a bad result (falsely indicating the result was good) in a mean number of 4.990 subjects (standard deviation; *SD* = 1.396) out of the 63 empirically good results (.079% false negative; 92.1% accuracy), and a false positive detection of a bad result (falsely indicating the result was bad) in a mean number of 6.990 subjects (*SD* = 1.179) out of the 77 empirically bad results (.091% false positive; 90.9% accuracy). When applying the trained model to the unrated subjects’ data, we identified 143 subjects with good results, 206 subjects with bad results, and 150 subjects with results selected for manual human rating. The manual human rating of the remaining subjects resulted in 10 more good results, for a total of 216. Out of these results, 197 had corresponding structural connectivity, demographic, and behavioral measures of interest and were therefore retained.

### Diffusion-weighted MRI

The dMRI imaging was performed using a twice-refocused sequence with a TR of 9100 ms, TE of 104 ms, FOV of 192×192 mm, and voxel size of 2×2×2 mm, with 30 directions of 66 axial slices having a b-value of 1000, 30 directions of 66 axial slices having a b-value of 2000, and 3 images of 66 axial slices having a b-value of 0. The structural connectivity (SC) measures of streamline probability and distance were calculated from the dMRI data using the TVB-UK Biobank pipeline (Frazier-Logue et al., 2022), which uses probabilistic tractography (*FSL bedpostx* to fit the probabilistic model and *probtrackx* to perform tractography; Hernandez-Fernandez et al., 2019; Jenkinson et al., 2012). The SC streamline probability is the number of connecting streamlines identified by the tractography divided by the total number of possible connections (i.e., normalized by the size of the region) and represents the probability of connection between all combinations of the 218 regions of interest in a combined atlas of the Schaefer 200 region atlas (Schaefer et al., 2018) and the subcortical Tian atlas (Tian et al., 2020). The subcortical regions were comprised of regions from the Tian Scale 1 atlas excluding the hippocampus. For hippocampus, the Scale 3 atlas was used with the two head divisions collapsed into a single parcel. The globus pallidus was excluded due to a large number of subjects without any detectible connections to or from this region, resulting in a total of 18 subcortical regions. The SC matrices were consistency thresholded (at least 50% of participants have the connection) and participants’ data were excluded if they did not have behavioral data, had regions with no connections, or had SC density (number of non-zero connections divided by the total number of possible connections) 3 SD or more away from the mean (retained *N* = 594).

### Visual Working Memory

Precision on the visual working memory task designed by Zhang & Luck (2008) was used to measure VWM (see Shafto et al., 2014 for more details). In this task 1 to 4 colored circles are presented peripherally to fixation and after a 900 ms delay participants are required to report the hue of the circle at the cued location. This measure declines significantly with age in this population, *R*(592) = −.291, *p* < .001 (see Figure 1; as demonstrated previously by Brockmole & Logie, 2013).

**Figure 1.**
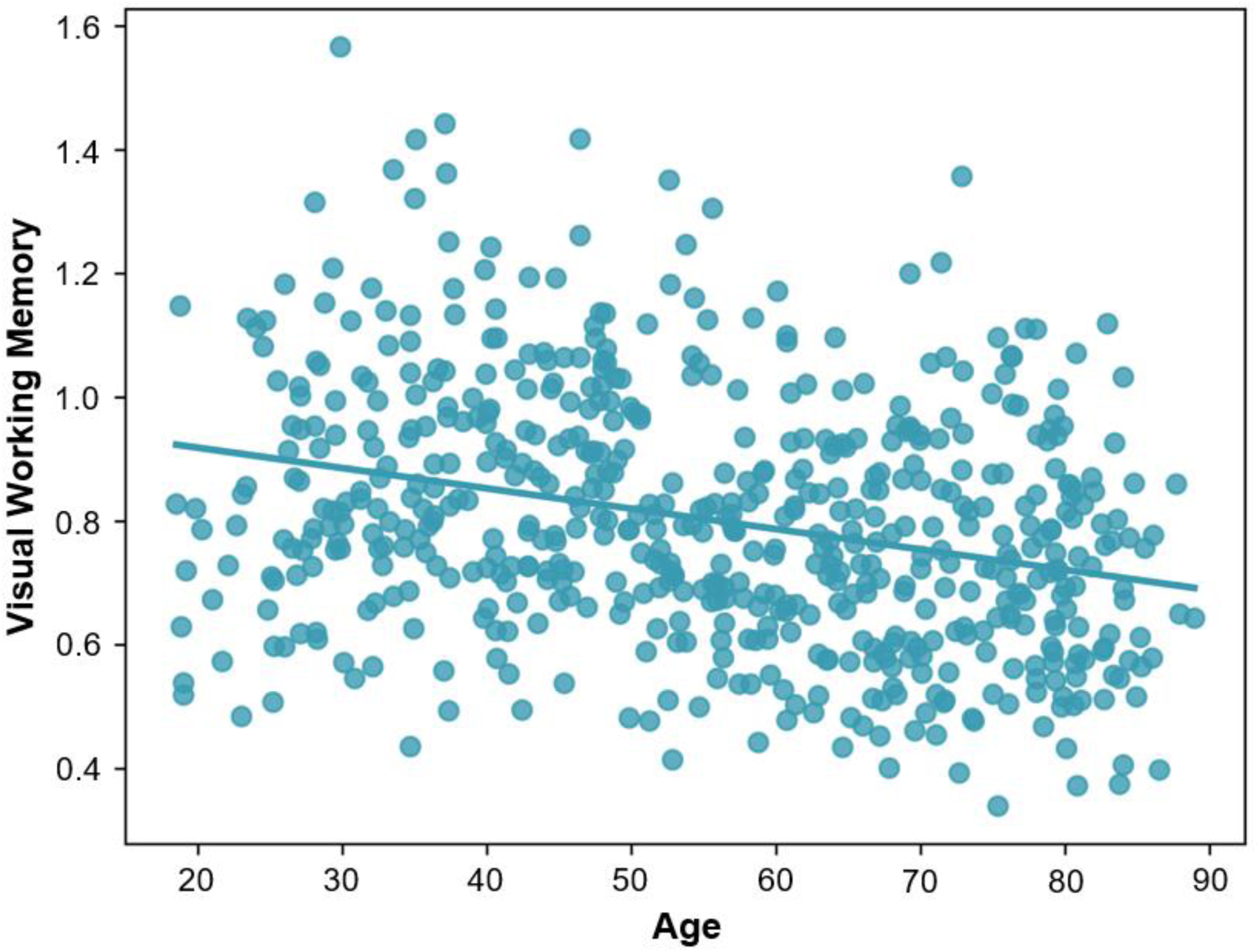
Relationship between age and visual working memory. Visual working memory decreases significantly with age, *R*(592) = −.291, *p* < .001.

### Dynamic Functional Connectivity

The rs-fMRI data was analyzed using a dynamic functional connectivity (dFC) analysis approach called Leading Eigenvector Dynamics Analysis (LEiDA; Cabral et al., 2017). This method uses phase coherence connectivity (e.g., Deco et al., 2017; Deco & Kringelbach, 2016; Glerean et al., 2012; Ponce-Alvarez et al., 2015) to compute a functional connectivity (FC) matrix at each timepoint of the resting-state fMRI scan. In contrast to other dFC methods that compare the full FC matrices across timepoints, LEiDA first computes the leading eigenvector of each FC matrix, making the method less susceptible to noise and better able to detect the recurrence of a particular state. The dFC matrices are separated into distinct states by applying a clustering analysis on the leading eigenvectors for all subjects and timepoints, resulting in states that are common to all subjects. The ideal number of states was chosen based on an evaluation of the clustering analysis that maximized the Dunn’s score (Dunn, 1973), average Sihouette coefficient (Rousseeuw, 1987), Calinski–Harabasz index (Caliński & Harabasz, 1974). With these states defined, each timepoint was then labelled according to which state the participant’s brain function was in at that timepoint, which allowed for calculation of the fractional occupancy (FO) of each state (probability of that state occurring at any given time) and the transition probability matrices (probability of the brain state changing from a specific state to another, or maintaining the same state, represented as a K×K matrix where K is the total number of states).

### Network Control Theory

Network control theory is a method that allows for observing the constraints that a structural connectivity network exerts on the functional dynamics of that system. By assuming a linear model of diffusion for simulating how activation spreads in parallel across the network, computationally efficient calculations can be performed to minimize the total input energy needed to guide the network from an initial state to a target state. The regions in the network that were given the largest amount of control energy were selected by this model as ideal recipients of the limited input energy. In the context of the brain, these control energy values provide insight into which regions are able to most efficiently drive a state transition given limited metabolic energy, and the utility of this method for network neuroscience has been well demonstrated in recent research (Gu et al., 2015, 2017; Kim & Bassett, 2020). This method provides an important opportunity to investigate how the structural connectivity network may reorganize to support a state transition identified as differing across the aging process via the LEiDA analysis of dynamic functional connectivity. The network control theory analysis was performed with *nctpy* (Parkes et al., 2023), using a continuous-time simulation, with a mixing parameter ρ of 1 to set equal weight to minimizing the control input energy and the neural activity, placing no constraints on which regions could be utilized for control inputs, and using a uniform full control set allowing all regions to act as controllers with equal control over the system dynamics.

### Partial Least Squares Analysis

Multivariate partial least squares (PLS) analysis (McIntosh & Lobaugh, 2004) was used to identify latent variables (LVs), each containing weights that describe the relationship of all brain measures with age and VWM. Behavioral PLS was used to examine effects of age and VWM as continuous measures. Models including sex as a group variable were examined, but produced no significant group effects so they were not included in the reported analyses. Secondary mean-centered PLS analyses were performed by categorizing subjects into 4 groups based on their age (YA or OA) and VWM (median split): YA with low VWM, YA with high VWM, OA with low VWM, and OA with high VWM. PLS uses singular value decomposition to project the data matrix onto orthogonal LVs (similar to canonical correlation analysis). The significance of the identified LVs was determined via permutation testing. We report only the most reliable PLS weights as determined by bootstrap resampling, which is used to calculate bootstrap ratios (BSR), which are the ratios of the PLS weights (saliences) to their standard errors as determined by bootstrap resampling (Kovacevic et al., 2013; McIntosh & Lobaugh, 2004). The resampling procedures were done using 1000 iterations for each. For the mean-centered PLS, brainscores were derived for each LV using the dot-product of the PLS weights for the brain metric and the values for the metric for each participant. The brainscores are similar to factor scores and can indicate the degree to which a subject or group show the pattern captured in the LV. In the present paper, we use these to convey the relative difference in LV expression between groups, using the mean brainscore and the bootstrap estimated standard error.

## Results

### Dynamic Functional Connectivity

LEiDA identified 5 states that were occupied during the course of the resting-state fMRI scan. The first state identified was the Global Coherence state, which is described as a state in which brain regions demonstrate overall coherence in their signals (Cabral et al., 2017). The second state overlapped primarily with a set of regions commonly referred to as the Default Mode Network (DMN). The third state overlapped primarily with the ventral attention network but also included dorsal regions from the dorsal attention network and will, therefore, be referred to simply as the Attention Network state. The fourth state overlapped primarily with the Frontoparietal Network (FPN), and the fifth state overlapped with regions corresponding to both the Somatomotor Network and the Visual Network (as defined by the Yeo et al., 2011 functional atlas; see Figure 2), and will be referred to as the Somatomotor/Visual Network state.

**Figure 2.**
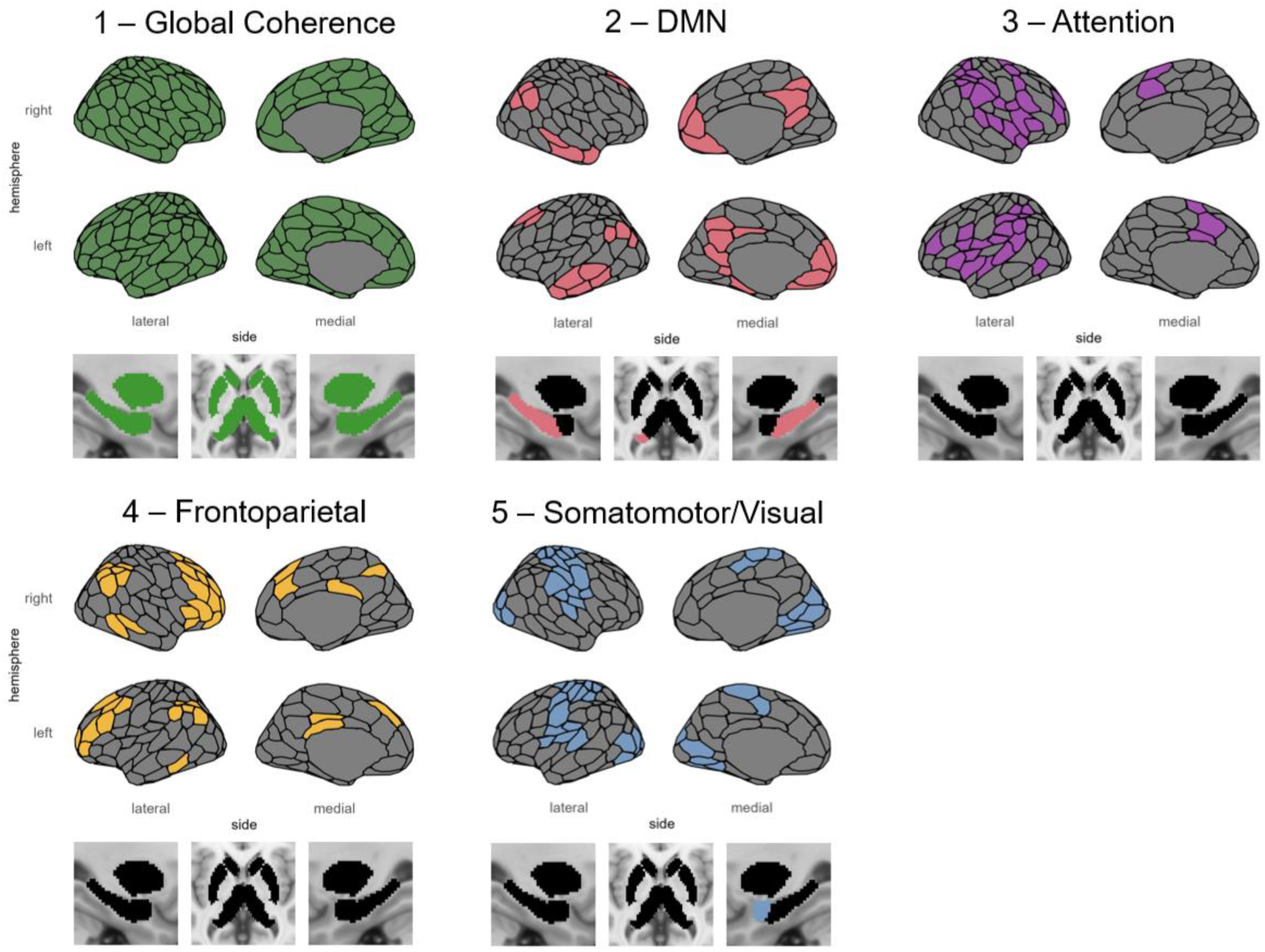
Dynamic functional connectivity states identified by the LEiDA analysis. State 1 in green represents the Global Coherence state, State 2 in pink represents the Default Mode Network (DMN) state, State 3 in purple represents the Attention Network state, State 4 in yellow represents the Frontoparietal Network (FPN) state, and State 5 in blue represents the Somatomotor/Visual Network state.

With these states defined, state timecourses were then produced for each individual, which allowed for the calculation of each individual’s fractional occupancy (FO; the percentage of time spent in each state). Behavioral PLS analyses were conducted with FO of each state as independent variables and age as the dependent variable in the first analysis, with VWM as the dependent variable in the second analysis, and with age and VWM as dependent variables in the third analysis. The first analysis with age identified a significant LV (permutation *p* = .048) that was positively associated with age, indicating that older adults spend more time in the DMN and FPN states (see Figure 3 A and B). The second analysis identified a significant LV (permutation *p* = .028) that was positively associated with VWM, indicating that less time spent in the Global Coherence state and more time spent in DMN was associated with better VWM (see Figure 3 C and D). The third analysis identified a significant LV (permutation *p* < .001) that was positively associated with both age and VWM, indicating that less time spent in the Global Coherence state and more time spent in DMN and FPN was associated with a better VWM in older age (see Figure 3 E and F). An additional mean-centered PLS was performed comparing FO in 4 groups: YA with low VWM, YA with high VWM, OA with low VWM, and OA with high VWM. This analysis identified 1 significant LV (permutation *p* < .001) with brainscores indicating that it identified how FO was associated with the OA high VWM group (YA low VWM: *brainscore* [95% CI] = −.0482 [−.079 to −.017]; YA high VWM: *brainscore* [95% CI] = −.007 [−.035 to .021]; OA low VWM: *brainscore* [95% CI] = −.022 [−.050 to .006]; OA high VWM: *brainscore* [95% CI] = .076 [.039 to .114]). This effect for OA with high VWM were consistent with the behavioral PLS LV associated with increasing age and VWM (see Figure 3 F), with BSRs that were reliably negative for Global Coherence and reliably positive for DMN and FPN states. The identified positive BSRs for DMN and FPN indicate that the increased time spent in DMN and FPN is not a detrimental aspect of the aging brain’s functional dynamics, but rather that this difference is advantageous for VWM.

**Figure 3.**
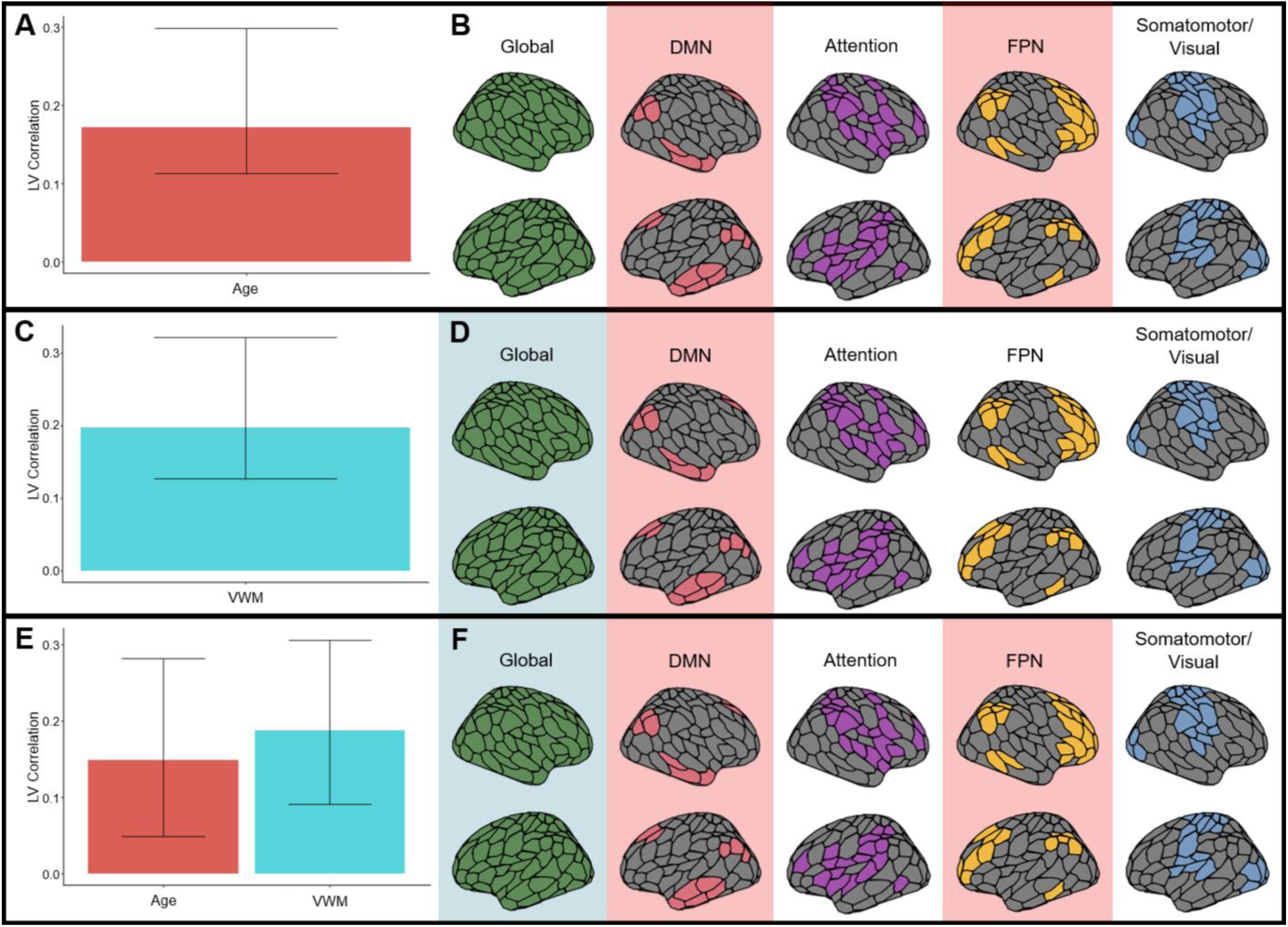
PLS analyses with independent variables of FO and dependent variable age (**A** and **B**), dependent variable VWM (**C** and **D**) and dependent variables of age and VWM (**E** and **F**). (**A**), (**C**), and (**E**) depict the behavioral correlation between the dependent variables and the LV, while (**B**), (**D**), and (**F**) represent the BSRs highlighting the reliable positive behavior-FO associations in red and the reliable negative behavior-FO associations in blue.

LEiDA also allows for the calculation of a state transition probability matrix, representing the probability of a transition occurring from state *i* in row *i* to state *j* in column *j* in the 5×5 matrix, where 5 is the total number of states (see Figure 4 B). Diagonal elements in the transition probability matrix represent maintenance of that state without a transition to another state. A PLS analysis with the transition probability matrix values as the independent variables and age as the dependent variable identified a significant LV (permutation *p* = .009). This LV was positively associated with age (see Figure 4 A), so that positive BSRs represent transitions (off diagonal elements) or maintenances (diagonal elements) that were more probable in older adults while negative BSRs represent transitions or maintenances that were less probable in older adults. The analysis identified the maintenance of State 2 (DMN) and State 3 (Attention) as being more prevalent in older adults, consistent with the FO analysis finding that DMN and FPN FO were greater for older adults, while adding Attention as another state that is more prevalent in older adults. The probability of transition from State 5 (Somatomotor/Visual) to State 3 (Attention) was also greater for older adults. Conversely, the probability of transition from State 2 (DMN) to State 1 (Global Coherence), from State 1 (Global Coherence) to State 4 (FPN), and from State 3 (Attention) to State 4 (FPN) were decreased in older age (see Figure 4 B as well as Figure 4 C for a graph representation of reliably age-related transitions and maintenances). As the only reliably age-related transition probability that increased with age, the transition from State 5 (Somatomotor/Visual) to State 3 (Attention) was investigated further from a structural connectivity perspective using network control theory.

**Figure 4.**
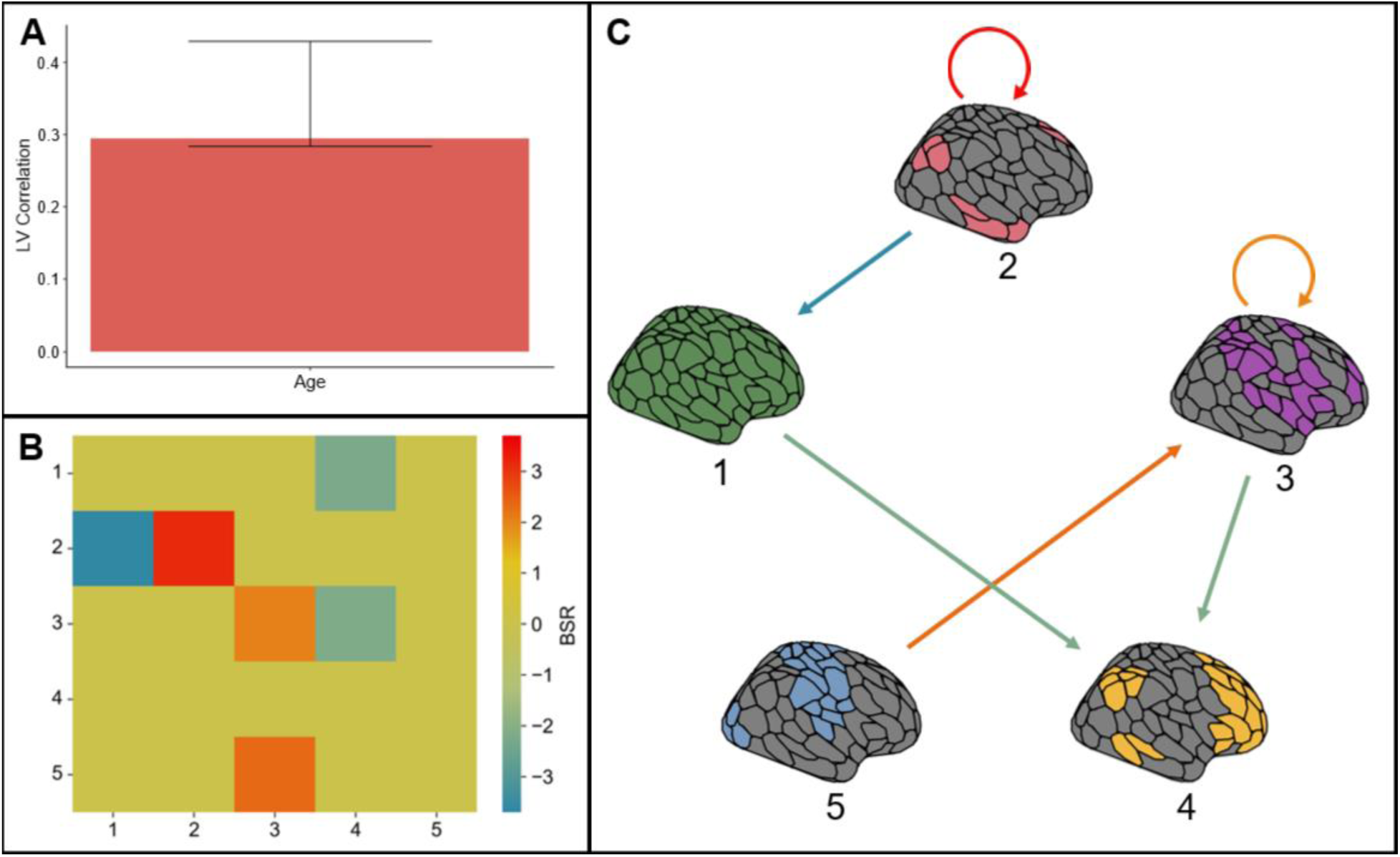
PLS analysis with independent variables of transition probabilities and dependent variable age. (**A**) depicts the behavioral correlation between age and the LV, while (**B**) and (**C**) represent the BSRs highlighting the reliable positive associations with age in red and the reliable negative associations with age in blue. BSRs are represented as both a transition probability matrix (**B**) and a transition probability graph (**C**).

### Network Control Theory

The network control theory analysis identified a combination of control energies given to each region corresponding to the optimal (minimized) total inputs to the simulation resulting in a transition from State 5 (Somatomotor/Visual) to State 3 (Attention; see Figure 5 A). A high value of control energy in a region can therefore be interpreted as that region being situated in the structural network architecture in a such a way that affords an ideal level of efficient control over the transition from the Somatomotor/Visual Network state to the Attention Network state (i.e., the brain relies heavily on that region for this transition). Separate PLS analyses for YA and OA were conducted, with control energies as independent variables and age and VWM as dependent variables. The YA analysis identified a significant LV (permutation *p* = .029) positively associated with age and VWM, indicating that VWM was better with age in individuals with greater control energies in regions with positive BSRs and with lower control energies in regions with negative BSRs (see Figure 5 B). The OA analysis identified a significant LV (permutation *p* < .001) negatively associated with age and positively associated with VWM, indicating that VWM was greater for individuals on the younger end of this age range if they had greater control energies in regions with positive BSRs and with lower control energies in regions with negative BSRs (see Figure 5 D). The YA and OA analyses both identified LVs associated with better VWM, and both analyses identified an effect of age towards the age category boundary of 50 years, indicating an effect on VWM that is maximized in middle age. The regions identified in these age groups had some overlap but also clear differences. For both YA and OA the RH hippocampus tail, RH caudate, LH central sulcus, LH frontal operculum, and bilateral dorsal parietal regions were negatively associated with VWM, indicating that a reliance on these regions for the Somatomotor/Visual to Attention state transition was disadvantageous (see Figure 5 C and E). Regions unique to YA demonstrating this negative relationship between control energy and VWM included RH inferior extrastriate, RH anterior insula, and LH anterior inferior temporal lobe (see Figure 5 C). Regions unique to OA demonstrating this negative relationship between control energy and VWM included the LH anterior thalamus, LH caudate, LH ventral prefrontal cortex (PFC), RH calcarine and RH striate cortex (see Figure 5 E). For the YA, higher control energies were associated with better VWM in a cluster of regions the RH insula and frontal operculum, as well as a RH dorsal parietal region, LH dorsolateral and medial PFC, LH inferior parietal lobule (IPL), and LH posterior cingulate cortex (PCC; see Figure 5 C). For the OA, higher control energies were associated with better VWM in two regions of the LH insula, LH postcentral gyrus, a cluster of regions in the LH posterior parietal cortex including the IPL, SPL, and postcentral gyrus, LH striate/calcarine, LH midcingulate, RH postcentral gyrus, RH medial parietal, and RH superior extrastriate (see Figure 5 E).

**Figure 5.**
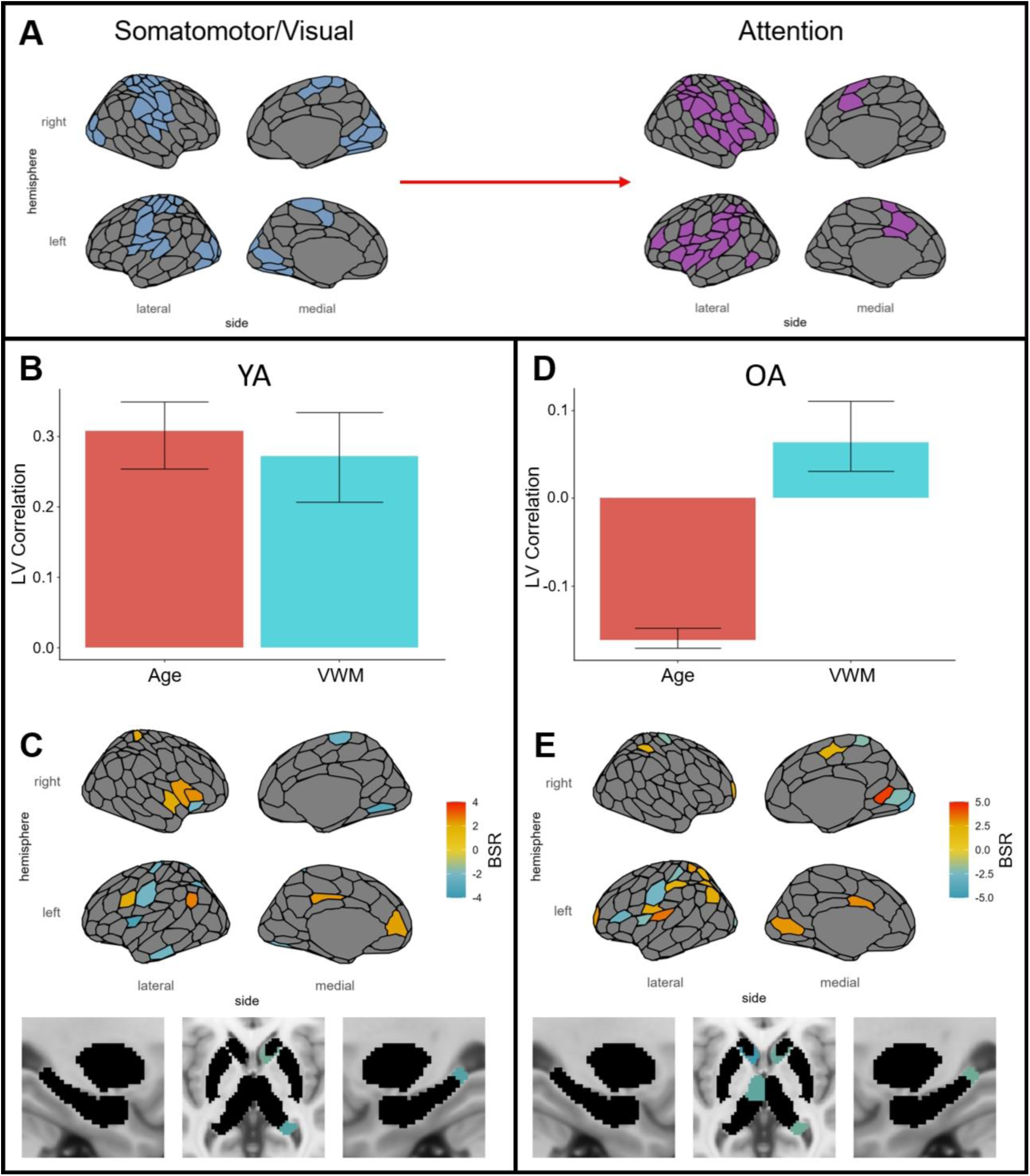
PLS analyses with independent variables of control energies for a state transition from the Somatomotor/Visual to the Attention state (**A**) and dependent variables of age and VWM. (**B**) and (**D**) depict the behavioral correlation of age and VWM with the LV, while (**C**) and (**E**) represent the BSRs highlighting the regions with reliable positive associations with age and VWM in yellow to red and the reliable negative associations with age and VWM in green to blue.

The secondary mean-centered PLS analysis identified 2 significant LVs. The first LV (permutation *p* < .001) identified an age contrast, whereby positive BSRs were associated with OA and negative BSRs were associated with YA (YA low VWM: *brainscore* [95% CI] = −.466 [−.588 to −.345]; YA high VWM: *brainscore* [95% CI] = −.119 [−.234 to −.004]; OA low VWM: *brainscore* [95% CI] = .188 [.075 to .300]; OA high VWM: *brainscore* [95% CI] = .398 [.287 to .508]; see Figure 6 A). The second LV (permutation *p* = .024) identified a VWM contrast within the YAs, whereby positive BSRs were associated with YA high VWM and negative BSRs were associated with YA low VWM (YA low VWM: *brainscore* [95% CI] = −.234 [−.370 to −.098]; YA high VWM: *brainscore* [95% CI] = .434 [.338 to .529]; OA low VWM: *brainscore* [95% CI] = −.104 [−.214 to .006]; OA high VWM: *brainscore* [95% CI] = −.096 [−.230 to .039]; see Figure 6 B).

**Figure 6.**
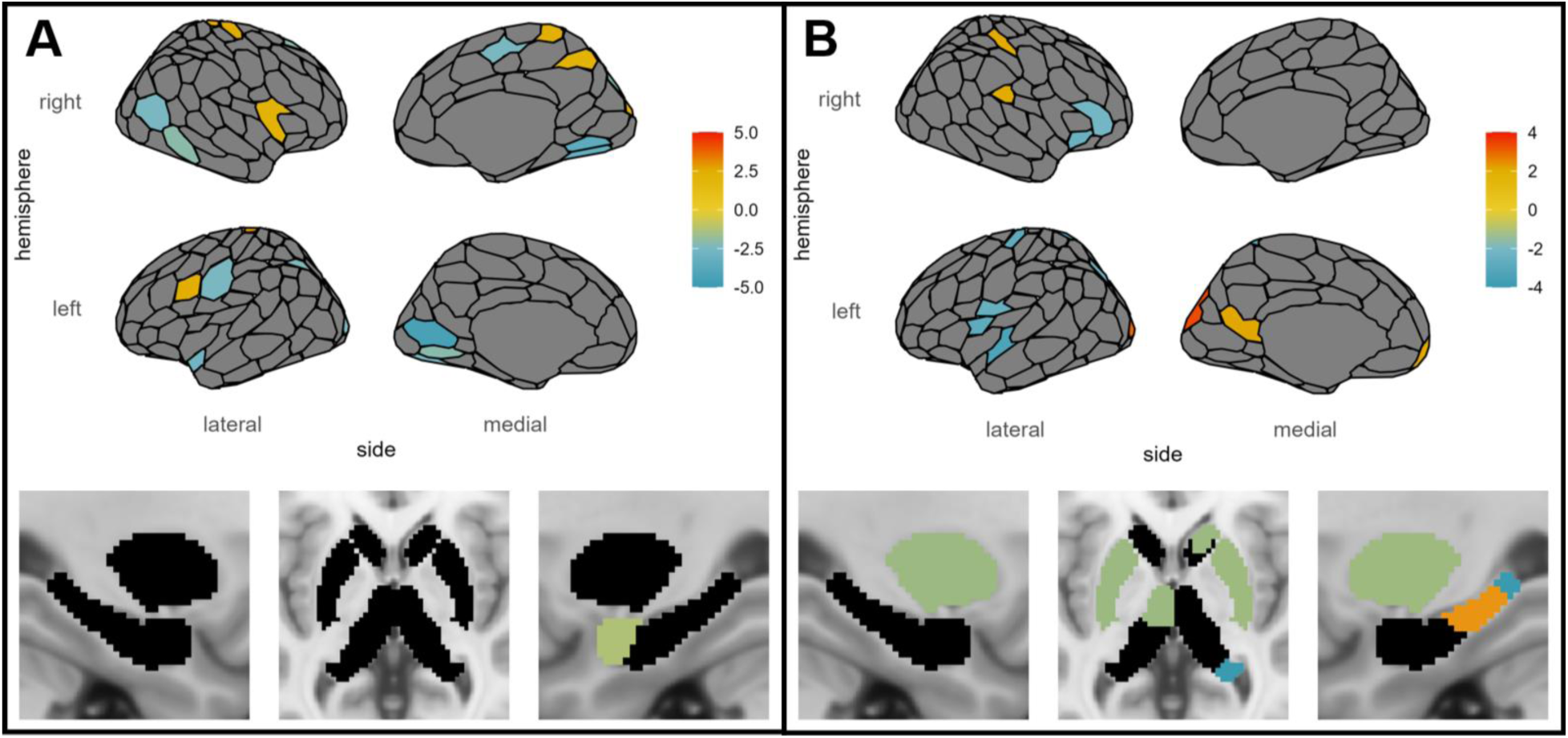
BSRs from mean-centered PLS, identifying 2 significant LVs. LV1 (**A**) identified an age contrast whereby positive BSRs are associated with OA and negative BSRs are associated with YA, while LV2 (**B**) identified a VWM contrast within the YA group, whereby positive BSRs are associated with high VWM and negative BSRs are associated with low VWM.

## Discussion

We have demonstrated that dynamic characteristics of the brain’s structural connectivity are related to data-driven measures of functional dynamics, that this relationship matures across the lifespan, and that elements of these dynamics support better VWM in middle age. Time spent in the DMN and FPN states increased with age and was associated with better VWM, as were decreases in Global Coherence time. Maintenance of the DMN and Attention Network states increased with age, while state transition probability from the DMN state to Global Coherence, Global Coherence to the FPN state, and the Attention Network state to the FPN state decreased with age. State transition probability from the Somatomotor/Visual Network state to the Attention Network state increased with age, and adaptations of the structural connectivity network supporting this state transition were associated with better VWM, particularly in middle age.

Our results highlight the importance of investigating *both* functional and structural brain networks from the perspective of how they support dynamics across the lifespan. Brain structure may be relatively fixed at physical and temporal macroscales, but the functions they support are dynamic. The dynamics that support better VWM with age provide an example of how the structural network informs the behavior of functional networks that support cognitive abilities. It is also interesting to see that VWM was associated with signatures of brain function during rest, separate from any related or demanding task, suggesting that individuals’ patterns of functional switching in the brain and the structural connections that make the transition between the Somatomotor/Visual Network state and the Attention Network state easier represent an important indicator of visual working memory as a person ages. The identification of the transition between these states specifically suggests that regardless of task demands (even at rest in this case), an individual’s ability to transition from a sensory/visual input state to an attention state that can attend to these inputs translates to their ability in a visual working memory task that requires taking visual inputs and attending to them effectively.

It is interesting to note that the structural connectivity network control theory analysis for younger adults implicated a number of regions associated positively with VWM that overlap with the Somatomotor/Visual and Attention networks, indicating that part of the role of influencing the efficiency of these transitions lies within these networks themselves, while also implicating regions from both the DMN and FPN. This suggests that performing the state transition efficiently relies on multinetwork coordination, and that this develops towards middle age. Conversely, regions positively associated with VWM for older adults were primarily within the Somatomotor/Visual and Attention networks, suggesting an ideal multinetwork coordination is established by middle age and that age-related differences occur locally within the Somatomotor/Visual and Attention networks after middle age.

### Connections to Past Research

The regions positively related to VWM in the network control theory analysis demonstrated consistency with past research, identifying the insula (RH for YA and LH for OA) as highlighted in previous network control theory research on WM task-based fMRI data (Cai et al., 2021). Furthermore, the dynamic functional connectivity analyses identified that time spent in the DMN and FPN states not was only not only greater in older adults, but that time spent in these states was associated with better VWM, adding to evidence from past research finding that the DMN becomes more segregated after WM training (Finc et al., 2020). Our finding that increased time spent in the FPN state at rest was associated with better VWM is also consistent with long-standing knowledge of the importance of the FPN to WM (D’Esposito, 2007; Smith & Jonides, 1999; Wager & Smith, 2003). The Global Coherence state identified by the LEiDA analysis represents a state in which brain regions are working together overall, rather than isolating a certain specialized subnetwork. Past research using this method has identified the Global Coherence state as being more prominently represented in older adults with good cognitive performance compared to those with poor cognitive performance (Cabral et al., 2017). Cognitive performance in this previous research was assessed using multiple measures sensitive to executive function, memory, and mood. Interestingly, our results found that decreases in the prevalence of Global Coherence were associated with better VWM. It seems the specific domain of cognitive ability may affect the relevance of time spent in the Global Coherence state, and that, for VWM, individuals with the ability to spend more time in specific states (DMN and FPN) rather than in a Global Coherence state, and to efficiently transition from the Sensorimotor/Visual to the Attention state, are more likely to have better VWM.

### Limitations and Future Directions

A longitudinal investigation utilizing this approach would provide a fuller picture of the relationship between structure, functional network dynamics, and cognitive ability. Recent research has highlighted the important differences between longitudinal and cross-sectional brain age research (Vidal-Pineiro et al., 2021). Adding a longitudinal perspective would represent an important contribution to this research investigating how the dynamic functional repertoires available are also dynamic across the lifespan, from infancy to childhood, adolescence, young adulthood, and old age. We know from both this and past research (Neudorf et al., 2024) that the network regimes that are ideal for healthy young adults are not the same as those that are ideal for older adults, so a continued exploration of brain networks across the lifespan with an eye for how these architectures address unique demands represents an important effort to diversify our understanding of how dynamic the brain network is across the lifespan.

## Conclusion

An investigation of the functional and structural brain networks’ dynamics allowed us to uncover how these different perspectives on brain dynamics are related. Furthermore, visual working memory relies on specifically tuned brain networks in middle age that can efficiently transition from a sensorimotor/visual focused state to a state wherein the individual is ready to attend to important information and maintain default mode and frontoparietal network states. The sensorimotor/visual to attention functional state transition was supported by the structural connectivity network, and different regions of the structural network were important during the approach to and the departure from middle age. This work has identified brain network signatures related to task performance based on brain structure and function at rest, suggesting that intrinsic features of brain dynamics may be valuable for assessing brain aging trajectories and relating to clinical conditions such as dementia.

## Acknowledgements

This research was supported by the Natural Sciences and Engineering Research Council of Canada (NSERC) through Postdoctoral Fellowships Program funding to Josh Neudorf, and by NSERC Discovery Grant *RGPIN-2018-04457* and Canadian Institutes of Health Research (CIHR) Project Grant *PJT-168980* to the senior author Anthony R. McIntosh. We thank Jaspreet Dodd for her assistance in validating the machine learning ratings of the fMRI data. This research was enabled in part by support provided by the British Columbia DRI Group and the Digital Research Alliance of Canada (alliancecan.ca). The authors affirm that there are no conflicts of interest to disclose.

## References

Bassett, D. S., Yang, M., Wymbs, N. F., & Grafton, S. T. (2015). Learning-induced autonomy of sensorimotor systems. Nature Neuroscience, 18(5), 744–751. 10.1038/nn.3993

Benkarim, O., Paquola, C., Park, B., Royer, J., Rodríguez-Cruces, R., Vos de Wael, R., Misic, B., Piella, G., & Bernhardt, B. C. (2022). A Riemannian approach to predicting brain function from the structural connectome. NeuroImage, 257, 119299. 10.1016/j.neuroimage.2022.119299

Brockmole, J. R., & Logie, R. H. (2013). Age-Related Change in Visual Working Memory: A Study of 55,753 Participants Aged 8–75. Frontiers in Psychology, 4. 10.3389/fpsyg.2013.00012

Cabral, J., Vidaurre, D., Marques, P., Magalhães, R., Silva Moreira, P., Miguel Soares, J., Deco, G., Sousa, N., & Kringelbach, M. L. (2017). Cognitive performance in healthy older adults relates to spontaneous switching between states of functional connectivity during rest. Scientific Reports, 7(1), Article 1. 10.1038/s41598-017-05425-7

Cai, W., Ryali, S., Pasumarthy, R., Talasila, V., & Menon, V. (2021). Dynamic causal brain circuits during working memory and their functional controllability. Nature Communications, 12(1), 3314. 10.1038/s41467-021-23509-x

Caliński, T., & Harabasz, J. (1974). A dendrite method for cluster analysis. Communications in Statistics, 3(1), 1–27. 10.1080/03610927408827101

Deco, G., Cabral, J., Woolrich, M. W., Stevner, A. B. A., van Hartevelt, T. J., & Kringelbach, M. L. (2017). Single or multiple frequency generators in on-going brain activity: A mechanistic whole-brain model of empirical MEG data. NeuroImage, 152, 538–550. 10.1016/j.neuroimage.2017.03.023

Deco, G., Jirsa, V. K., & McIntosh, A. R. (2011). Emerging concepts for the dynamical organization of resting-state activity in the brain. Nature Reviews Neuroscience, 12(1), 43–56. 10.1038/nrn2961

Deco, G., & Kringelbach, M. L. (2016). Metastability and Coherence: Extending the Communication through Coherence Hypothesis Using A Whole-Brain Computational Perspective. Trends in Neurosciences, 39(3), Article 3. 10.1016/j.tins.2016.01.001

D’Esposito, M. (2007). From cognitive to neural models of working memory. Philosophical Transactions of the Royal Society B: Biological Sciences, 362(1481), 761–772. 10.1098/rstb.2007.2086

Dunn, J. C. (1973). A Fuzzy Relative of the ISODATA Process and Its Use in Detecting Compact Well-Separated Clusters. Journal of Cybernetics, 3(3), 32–57. 10.1080/01969727308546046

Faber, S. E. M., Belden, A. G., Loui, P., & McIntosh, R. (2023). Age-related variability in network engagement during music listening. Network Neuroscience, 7(4), 1404–1419. 10.1162/netn_a_00333

Faber, S. E., McIntosh, A. R., Brown, T., & Carpentier, S. (2024). Mapping multi-modal dynamic network activity during naturalistic music listening (p. 2023.07.05.547865). bioRxiv. 10.1101/2023.07.05.547865

Finc, K., Bonna, K., He, X., Lydon-Staley, D. M., Kühn, S., Duch, W., & Bassett, D. S. (2020). Dynamic reconfiguration of functional brain networks during working memory training. Nature Communications, 11(1), 2435. 10.1038/s41467-020-15631-z

Frazier-Logue, N., Wang, J., Wang, Z., Sodums, D., Khosla, A., Samson, A. D., McIntosh, A. R., & Shen, K. (2022). A Robust Modular Automated Neuroimaging Pipeline for Model Inputs to TheVirtualBrain. Frontiers in Neuroinformatics, 16, 883223. 10.3389/fninf.2022.883223

Glerean, E., Salmi, J., Lahnakoski, J. M., Jääskeläinen, I. P., & Sams, M. (2012). Functional Magnetic Resonance Imaging Phase Synchronization as a Measure of Dynamic Functional Connectivity. Brain Connectivity, 2(2), 91–101. 10.1089/brain.2011.0068

Gu, S., Betzel, R. F., Mattar, M. G., Cieslak, M., Delio, P. R., Grafton, S. T., Pasqualetti, F., & Bassett, D. S. (2017). Optimal trajectories of brain state transitions. NeuroImage, 148, 305–317. 10.1016/j.neuroimage.2017.01.003

Gu, S., Pasqualetti, F., Cieslak, M., Telesford, Q. K., Yu, A. B., Kahn, A. E., Medaglia, J. D., Vettel, J. M., Miller, M. B., Grafton, S. T., & Bassett, D. S. (2015). Controllability of structural brain networks. Nature Communications, 6(1), 8414. 10.1038/ncomms9414

Harrison, S. A., & Tong, F. (2009). Decoding reveals the contents of visual working memory in early visual areas. Nature, 458(7238), Article 7238. 10.1038/nature07832

Haussler, D., Krogh, A., Mian, S., & Sjolander, K. (1992). PROTEIN MODELING USING HIDDEN MARKOV MODELS: ANALYSIS OF GLOBINS. University of California at Santa Cruz.

Heisz, J. J., Gould, M., & McIntosh, A. R. (2015). Age-related Shift in Neural Complexity Related to Task Performance and Physical Activity. Journal of Cognitive Neuroscience, 27(3), Article 3. 10.1162/jocn_a_00725

Hernandez-Fernandez, M., Reguly, I., Jbabdi, S., Giles, M., Smith, S., & Sotiropoulos, S. N. (2019). Using GPUs to accelerate computational diffusion MRI: From microstructure estimation to tractography and connectomes. NeuroImage, 188, 598–615. 10.1016/j.neuroimage.2018.12.015

Honey, C. J., Sporns, O., Cammoun, L., Gigandet, X., Thiran, J. P., Meuli, R., & Hagmann, P. (2009). Predicting human resting-state functional connectivity from structural connectivity. Proceedings of the National Academy of Sciences, 106(6), Article 6. 10.1073/pnas.0811168106

Jenkinson, M., Beckmann, C. F., Behrens, T. E. J., Woolrich, M. W., & Smith, S. M. (2012). FSL. NeuroImage, 62(2), Article 2. 10.1016/j.neuroimage.2011.09.015

Kim, J. Z., & Bassett, D. S. (2020). Linear Dynamics and Control of Brain Networks. In B. He (Ed.), Neural Engineering (pp. 497–518). Springer International Publishing. 10.1007/978-3-030-43395-6_17

Kovacevic, N., Abdi, H., Beaton, D., & McIntosh, A. R. (2013). Revisiting PLS Resampling: Comparing Significance Versus Reliability Across Range of Simulations. In H. Abdi, W. W. Chin, V. Esposito Vinzi, G. Russolillo, & L. Trinchera (Eds.), New Perspectives in Partial Least Squares and Related Methods (pp. 159–170). Springer. 10.1007/978-1-4614-8283-3_10

Littlejohns, T. J., Holliday, J., Gibson, L. M., Garratt, S., Oesingmann, N., Alfaro-Almagro, F., Bell, J. D., Boultwood, C., Collins, R., Conroy, M. C., Crabtree, N., Doherty, N., Frangi, A. F., Harvey, N. C., Leeson, P., Miller, K. L., Neubauer, S., Petersen, S. E., Sellors, J., … Allen, N. E. (2020). The UK Biobank imaging enhancement of 100,000 participants: Rationale, data collection, management and future directions. Nature Communications, 11(1), Article 1. 10.1038/s41467-020-15948-9

Lynn, C. W., & Bassett, D. S. (2019). The physics of brain network structure, function and control. Nature Reviews Physics, 1(5), 318–332. 10.1038/s42254-019-0040-8

McIntosh, A. R., & Lobaugh, N. J. (2004). Partial least squares analysis of neuroimaging data: Applications and advances. NeuroImage, 23 Suppl 1, S250–263. 10.1016/j.neuroimage.2004.07.020

Neudorf, J., Kress, S., & Borowsky, R. (2022). Structure can predict function in the human brain: A graph neural network deep learning model of functional connectivity and centrality based on structural connectivity. Brain Structure and Function, 227(1), Article 1. 10.1007/s00429-021-02403-8

Neudorf, J., Shen, K., & McIntosh, A. R. (2024). Reorganization of Structural Connectivity in the Brain Supports Preservation of Cognitive Ability in Healthy Aging. Network Neuroscience, 1–49. 10.1162/netn_a_00377

Parkes, L., Kim, J. Z., Stiso, J., Brynildsen, J. K., Cieslak, M., Covitz, S., Gur, R. E., Gur, R. C., Pasqualetti, F., Shinohara, R. T., Zhou, D., Satterthwaite, T. D., & Bassett, D. S. (2023). Using network control theory to study the dynamics of the structural connectome (p. 2023.08.23.554519). bioRxiv. 10.1101/2023.08.23.554519

Ponce-Alvarez, A., Deco, G., Hagmann, P., Romani, G. L., Mantini, D., & Corbetta, M. (2015). Resting-State Temporal Synchronization Networks Emerge from Connectivity Topology and Heterogeneity. PLOS Computational Biology, 11(2), e1004100. 10.1371/journal.pcbi.1004100

Rousseeuw, P. J. (1987). Silhouettes: A graphical aid to the interpretation and validation of cluster analysis. Journal of Computational and Applied Mathematics, 20, 53–65. 10.1016/0377-0427(87)90125-7

Sarwar, T., Tian, Y., Yeo, B. T. T., Ramamohanarao, K., & Zalesky, A. (2021). Structure-function coupling in the human connectome: A machine learning approach. NeuroImage, 226, 117609. 10.1016/j.neuroimage.2020.117609

Schaefer, A., Kong, R., Gordon, E. M., Laumann, T. O., Zuo, X.-N., Holmes, A. J., Eickhoff, S. B., & Yeo, B. T. T. (2018). Local-Global Parcellation of the Human Cerebral Cortex from Intrinsic Functional Connectivity MRI. Cerebral Cortex, 28(9), 3095–3114. 10.1093/cercor/bhx179

Schirner, M., McIntosh, A. R., Jirsa, V., Deco, G., & Ritter, P. (2018). Inferring multi-scale neural mechanisms with brain network modelling. eLife, 7, e28927. 10.7554/eLife.28927

Schurgin, M. W. (2018). Visual memory, the long and the short of it: A review of visual working memory and long-term memory. *Attention, Perception*, & Psychophysics, 80(5), 1035– 1056. 10.3758/s13414-018-1522-y

Shafto, M. A., Tyler, L. K., Dixon, M., Taylor, J. R., Rowe, J. B., Cusack, R., Calder, A. J., Marslen-Wilson, W. D., Duncan, J., Dalgleish, T., Henson, R. N., Brayne, C., Matthews, F. E., & Cam-CAN. (2014). The Cambridge Centre for Ageing and Neuroscience (Cam-CAN) study protocol: A cross-sectional, lifespan, multidisciplinary examination of healthy cognitive ageing. BMC Neurology, 14(1), 204. 10.1186/s12883-014-0204-1

Smith, E. E., & Jonides, J. (1999). Storage and Executive Processes in the Frontal Lobes. Science, 283(5408), 1657–1661. 10.1126/science.283.5408.1657

Tian, Y., Margulies, D. S., Breakspear, M., & Zalesky, A. (2020). Topographic organization of the human subcortex unveiled with functional connectivity gradients. Nature Neuroscience, 23(11), Article 11. 10.1038/s41593-020-00711-6

Todd, J. J., & Marois, R. (2004). Capacity limit of visual short-term memory in human posterior parietal cortex. Nature, 428(6984), Article 6984. 10.1038/nature02466

Vidaurre, D., Smith, S. M., & Woolrich, M. W. (2017). Brain network dynamics are hierarchically organized in time. Proceedings of the National Academy of Sciences, 114(48), Article 48. 10.1073/pnas.1705120114

Wager, T. D., & Smith, E. E. (2003). Neuroimaging studies of working memory: *Cognitive, Affective*, & Behavioral Neuroscience, 3(4), 255–274. 10.3758/CABN.3.4.255

Yeo, B. T. T., Krienen, F. M., Sepulcre, J., Sabuncu, M. R., Lashkari, D., Hollinshead, M., Roffman, J. L., Smoller, J. W., Zöllei, L., Polimeni, J. R., Fischl, B., Liu, H., & Buckner, R. L. (2011). The organization of the human cerebral cortex estimated by intrinsic functional connectivity. Journal of Neurophysiology, 106(3), 1125–1165. 10.1152/jn.00338.2011

Zhang, W., & Luck, S. J. (2008). Discrete fixed-resolution representations in visual working memory. Nature, 453(7192), Article 7192. 10.1038/nature06860

Zimmermann, J., Ritter, P., Shen, K., Rothmeier, S., Schirner, M., & McIntosh, A. R. (2016). Structural architecture supports functional organization in the human aging brain at a regionwise and network level. Human Brain Mapping, 37(7), 2645–2661. 10.1002/hbm.23200

